# Endosomal structure and APP biology are not altered in preclinical cellular models of Down syndrome

**DOI:** 10.1101/2021.12.30.474570

**Authors:** Claudia Cannavo, Karen Cleverley, Cheryl Maduro, Paige Mumford, Dale Moulding, Elizabeth M. C. Fisher, Frances K. Wiseman

**Affiliations:** UK Dementia Research Institute at UCL, London, WC1N 3BG UK; Department of Neuromuscular Disease, UCL Queen Square Institute of Neurology, London, WC1N 3BG UK; Light Microscopy Core Facility, Great Ormond Street Institute of Child Health, University College, London, WC1N 1EH UK; LonDownS Consortium

## Abstract

Individuals who have Down syndrome (trisomy 21) are at greatly increased risk of developing Alzheimer’s disease – dementia. Alzheimer’s disease is characterised by the accumulation in the brain of amyloid-β plaques that are a product of amyloid precursor protein, encoded by the *APP* gene on chromosome 21. In Down syndrome the first site of amyloid-β accumulation is within endosomes and changes to endosome biology occur early in disease. Here we determine if primary mouse embryonic fibroblasts isolated from two mouse models of Down syndrome can be used to study endosome and APP cell biology. We report that in these cellular models of Down syndrome endosome number, size and APP processing are not altered, likely because *APP* is not dosage sensitive in these models, despite three copies of *App*.

## Introduction

Individuals with Down syndrome (DS), which is caused by trisomy of human chromosome 21 (Hsa21), have a high risk of developing early onset Alzheimer’s disease (AD). One of the earliest neuropathological features of AD in people who have DS is the intracellular accumulation of amyloid-β in the brain, followed by the accumulation of extracellular amyloid-β plaques [1]. Amyloid-β is a product of the *APP* gene that is encoded on Hsa21. Clinical-genetic studies indicate that three copies of *APP* are both sufficient and necessary for the development of early onset AD in people who have DS and in the general population. However, growing evidence suggests that other genes on Hsa21 can affect APP/amyloid-β, including via modulation of endosomal biology [2].

APP follows the central secretory pathway. Full-length APP is synthesised in the endoplasmic reticulum, transported to the Golgi and then to the plasma membrane [3,4]. From there, APP is internalized through endocytosis and either recycled to the plasma membrane or Golgi, or directed for degradation to the endo-lysosomes [5–7]. Where APP lies in the cell is important for its degradation. APP mainly undergoes two alternative types of processing, through the action of different secretases. The most common processing pathway is ‘non-amyloidogenic’, which principally occurs at the plasma membrane and consists of sequential cleavage by α- and γ-secretases. The second ‘amyloidogenic’ pathway leads to the production of amyloid-β, mainly occurs in endosomes and is mediated by sequential cleavage of APP by β- and γ-secretases [8,9]. Cleavage by β-secretase occurs first, and results in the production of an extracellular fragment that is released from the cell (sAPPβ) and of a transmembrane fragment (β-CTF) which is then cleaved by γ-secretase to produce amyloid-β.

Endosomal dysfunction and enlargement is observed in the brains of people who have AD and DS before amyloid-β plaque accumulation and has been suggested to a be a key factor in AD development [10–12]. Indeed this has been reported in early gestation of individuals with DS [11,13], in cells isolated from individuals with DS [14], in iPSCs-derived trisomy-21 neurons and organoids [15–17], and in mouse models of DS [18]. Whether this enlargement is caused by an increased fusion of endosomal bodies or an increase in the volume of single endosomes is disputed [19], likely because of the technical challenges encountered in the precise quantification of the very small endosomal bodies [20–22].

*APP* triplication is necessary for early endosomal dysfunction in DS models and is mediated by raised β-CTF [18,21]. Other Hsa21 genes/proteins may also contribute to this dysfunction [18,23]. For example, synaptojanin-1 (*SYNJ1*), is a phosphatase that mediates the uncoating of clathrin-coated vesicles. SYNJ1 levels are increased in the brains of people who have DS, and its overexpression causes endosomal enlargement [23]. The Hsa21 gene Intersectin-1 (*ITSN1*) encodes a regulator of endocytosis [24] and its levels are increased in DS [25]. Overexpression of the Regulator of Calcineurin 1 (*RCAN1*) affects vesicle recycling and endocytosis via its effect on calcineurin activity [26]. Finally, the Hsa21 microRNA gene *miR-155* negatively regulates the transcription of SNX27, a component of the retromer complex, and SNX27 levels are decreased in DS [27]. Since APP is subject to retrograde transport, impairment of this mechanism could lead to a longer residency of APP inside early endosomes, causing a change in early endosome structure, increased amyloidogenic processing of APP and modifying APP half-life [28].

In addition, research in preclinical systems suggests that genes on Hsa21 including *DYRK1A* and *BACE2* can modulate APP/Aβ biology when in three-copies [17,29,30]. Overexpression of DYRK1A in the brain of APP transgenic mice increases the total abundance of APP and Aβ via phosphorylation of APP at Thr668. BACE2 mostly functions as θ-secretase but may also degrade Aβ or cleave APP at the β-secretase site [31–33]. A recent study in organoids generated from trisomy 21 iPSCs demonstrated that three copies of *BACE2* protect against amyloid-β accumulation in that system [17]. These findings are consistent with gene-association studies implicating these genes in AD-risk in individuals who have DS [34–36].

Here, we investigate whether novel cellular models of DS that carry three copies of 114 or 30 mouse gene homologues of Hsa21 genes including *App, Synj1, Itsn1, Rcan1, Mir155, Dyrk1A* and *Bace2* can be used to study APP/amyloid-β and endosomal biology.

## Results

### Three copies of Hsa21 gene homologues in Dp1Tyb and Dp2Tyb mouse embryonic fibroblasts do not alter endosome numbers

We aimed to determine if an additional copy of Hsa21 homologues previously implicated in changed endosomal biology in DS were sufficient to increase the number or size of endosomes in mouse embryonic fibroblasts (MEFs) derived from segmental duplication mouse models of DS. Therefore, we established a systematic workflow for quantification of the number and the size distribution of early endosomes, using RAB5 staining, confocal imaging and deconvolution (**Supplementary Fig 1**). This workflow was validated by the over-expression of GFP-Rab5CA (Q79L) (RAB5CA), [37]) in wildtype (WT) MEFs (**Supplementary Fig 2**), leading to the expression of constitutively active RAB5 which enlarges endosomal bodies.

We then used this system to study MEFs derived from Dp(16Lipi-Dbtb21)1TybEmcf [herein referred to as Dp1Tyb] and Dp(16Mis18a-Runx1)2TybEmcf [Dp2Tyb] hemizygous mouse models of DS. The Dp1Tyb mouse has a segmental duplication of mouse chromosome 16 (Mmu16) that is homologous with Hsa21 and has an additional copy of 114 mouse orthologues of Hsa21 genes [38,39], including *App, Synj1, Itsn1, Rcan1, Mir155, Dyrk1A* and *Bace2*. The Dp2Tyb mouse model carries a segmental duplication of a smaller region of Mmu16 [38,39], of 30 genes including *Synj1* and *Itsn1* but does not contain an additional copy of *App or Mir155 (***Fig. 1A**).

**Figure 1.**
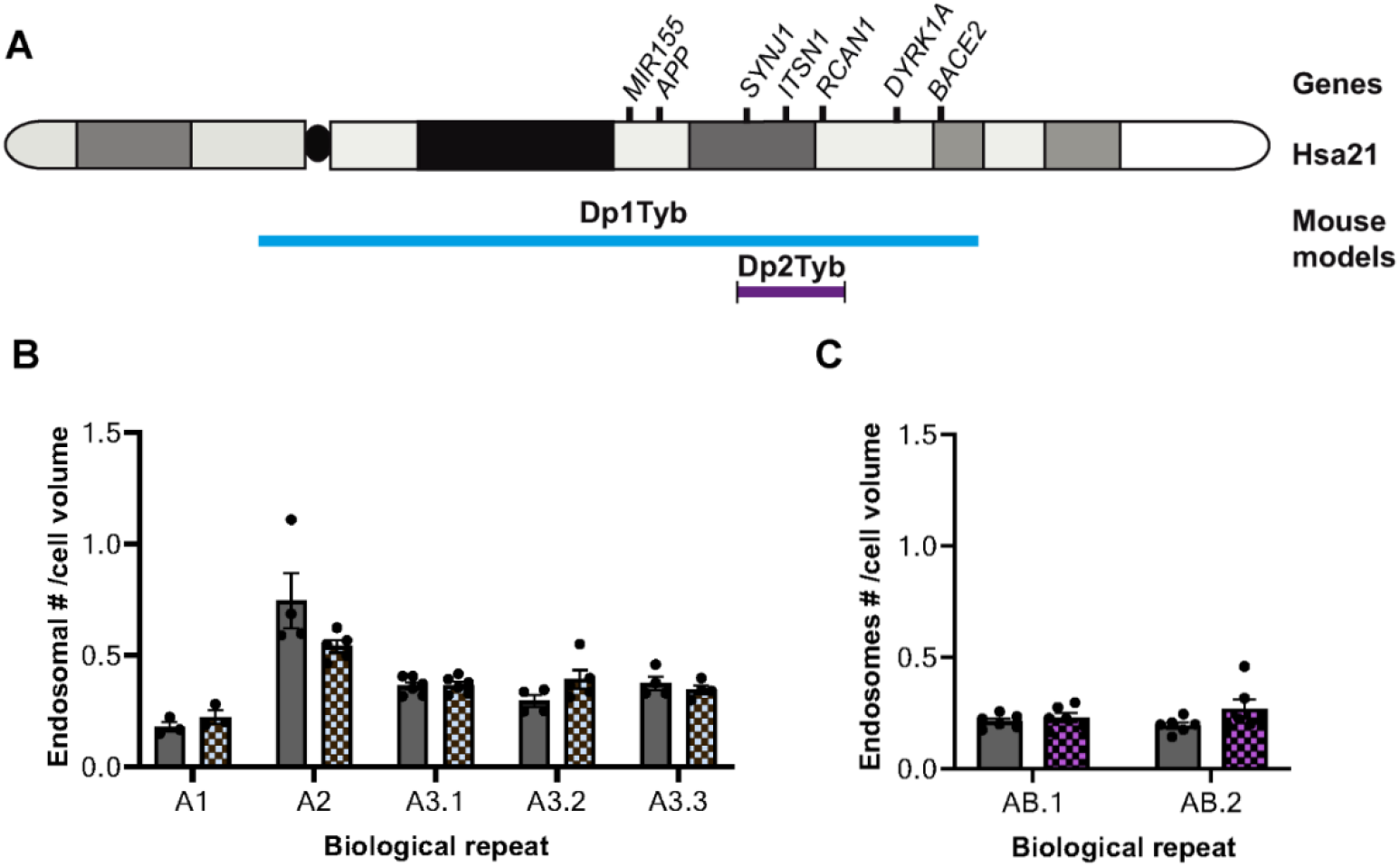
Number of endosomes per cell is not different in WT and Dp1Tyb or Dp2Tyb MEFs. **A)** Schematic of gene content of Dp1Tyb and Dp2Tyb mouse models **B)** No difference was found in the number of RAB5^+^ endosomes (normalised to cell volume) in WT and Dp1Tyb MEFs (Nested t-test *p* = 0.83, N = 5 biological repeats, N = 3-5 of technical repeats). **C**) No difference was found in the number of RAB5^+^ endosomes (normalised to cell volume) in WT and Dp2Tyb MEF (Nested t-test, *p* = 0.08, N = 2 biological repeats (independent MEF lines), N = 6 technical repeats). *Error bars = SEM*.

Using our workflow, we found no difference in the number of RAB5^+^ endosomes normalised to cell volume in WT and Dp1Tyb MEFs (WT = 2218 ± 58; Dp2Tyb = 2503 ± 119, N = 5 biological repeats) (**Fig. 1B**). Notably, biological variation in the number of RAB5^+^ endosomes between MEF isolates from individual litters of mice was observed, necessitating for our onward analysis a nested design that enabled us to compare the two genotypes while accounting for the variability between litters. Using the same methods described above, no difference was found in the number of endosomes in WT and Dp2Tyb MEFs (WT = 2809 ± 101; Dp2Tyb = 3306 ± 266 N = 2 biological replicates) (**Fig. 1C**).

### Three copies of Hsa21 mouse homologues in Dp1Tyb and Dp2Tyb MEFs do not increase endosomal volume

Using our workflow, we determined the volume distribution of RAB5^+^ endosomes in WT and Dp1Tyb MEFs. The average volume for the total number of endosomes was consistent with our initial pipeline studies in WT MEFs (WT = 0.07 ± 0.001 μm^3^, Dp1Tyb = 0.06 ± 0.001 μm^3^). WT MEFs isolated from the littermates of the segmental duplication mice were used to determine the 50 and 90 percentile values of endosomal volumes and these data were used to classify endosomes from both genotypes as small (0–50 percentile), medium (50–90 percentile) and large (> 90%). No difference in size distribution was found between WT and Dp1Tyb MEFs. The average volumes of endosomes classified as ‘large’ were compared and no difference was found between WT and Dp1Tyb MEFs (average volume of large endosomes: WT = 0.29 ± 0.006 μm^3^, Dp1Tyb = 0.27 ± 0.003 μm^3^ N = 5 biological repeats) (**Fig 2A, C**). Using the same method, no difference in the volume distribution or average volume of the large endosomes was found in WT and Dp2Tyb MEFs (average volume: WT = 0.96 ± μm^3^, Dp2Tyb = 0.71 ± 0.63 μm^3^ N = 2 biological repeats) (**Fig. 2B, D**). Two different confocal microscopes (LSM800 and LSM880) were used for the Dp1Tyb and Dp2Tyb studies, resulting in the difference in absolute WT endosomal volume across the two experiments.

**Figure 2.**
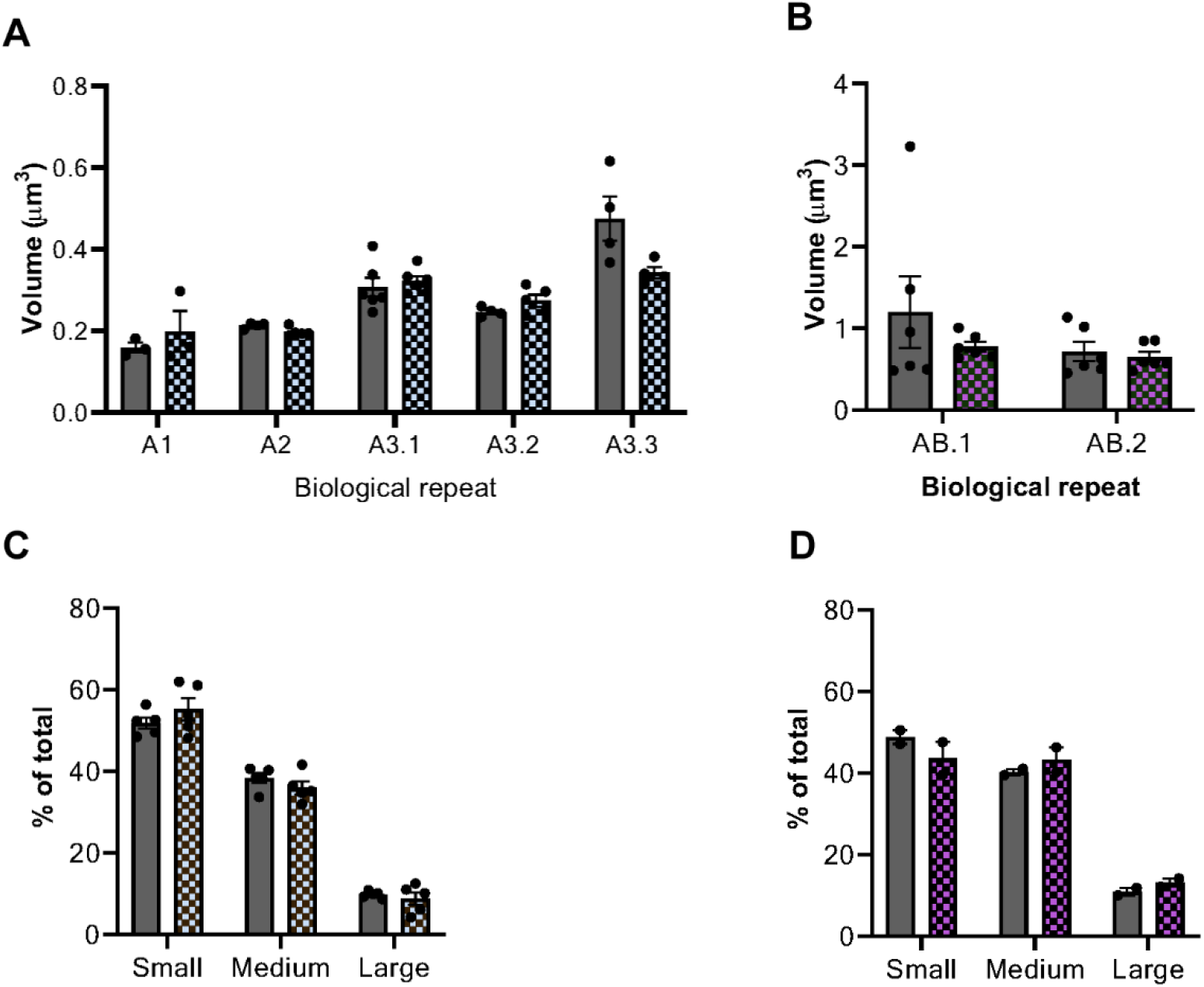
Endosomal volume distribution and mean volume of the largest endosomes are not different in WT and Dp1Tyb or Dp2Tyb MEFs. Endosomes were binned in three size categories: small (0-50 percentile of WT MEFs), medium (50-90 percentile of WT MEFs) and large (90-100 percentile of WT MEFs). The categories were determined using the endosomes in WT MEFs. **A)** No difference in RAB5^+^ endosome volume distribution was observed in WT and Dp1Tyb MEFs (Mann-Whitney U test). N = 5 biological repeats, N = 3-5 of technical repeats. **B)** No difference in RAB5^+^ endosome volume distribution was observed in WT and Dp2Tyb MEFs (Mann-Whitney U test). N = 2 biological repeats, N = 6 technical repeats. **C)** Volume of the RAB5^+^ endosomes classified as ‘large’ is not different in WT and Dp1Tyb MEFs (Nested t-test, *p* = 0.21). N = 5 of biological repeats (independent MEF lines), N = 3-5 of technical repeats. **D)** Volume of the RAB5^+^ endosomes classified as ‘large’ is not different in WT and Dp2Tyb MEFs (Nested t-test, *p* = 0.31 N = 2 biological repeats (independent MEF lines), N = 6 technical repeats). *Error bars = SEM*.

### Three-copies of *App* do not lead to raised APP protein level or altered half-life in the Dp1Tyb MEF model system

Previous work has suggested that three copies of *APP* and the resulting raised levels of APP protein and the APP cleavage product β-CTF are critical to the enlargement of early endosomes in the context of DS ([40]). Thus, we determined if three copies of *App* were sufficient to raise APP protein level in the Dp1Tyb MEFs or alter the protein half-life. We crossed the Dp1Tyb mouse model with a heterozygous *App* knockout animal *App*^*tm1Dbo*^ (*App*^*+/-*^) to generate MEFs and studied three of the resulting genotypes: Dp1Tyb with 3 copies of *App* (Dp1Tyb), Dp1Tyb/*App*^*+/*-^ with 2 copies of *App* and WT with 2 copies of *App*.

MEFs with the three genotypes were treated with cycloheximide and collected at 0 h, 15 min, 30 min, 1 h, 2 h and 4 h. APP protein abundance at each time point was measured by western blotting and a non-linear regression test was used to determine APP half-life. We found no difference in APP abundance or APP half-life in Dp1Tyb, Dp1Tyb/*App*^*+/*-^ and WT MEFs, suggesting that trisomy of Hsa21-homologous genes on Mmu16 including *App* is not sufficient to increase APP protein level in this cellular model and that this dosage-insensitivity is not the result of an increase in the proteins degradation rate (**Fig. 3 A-C**).

**Figure 3.**
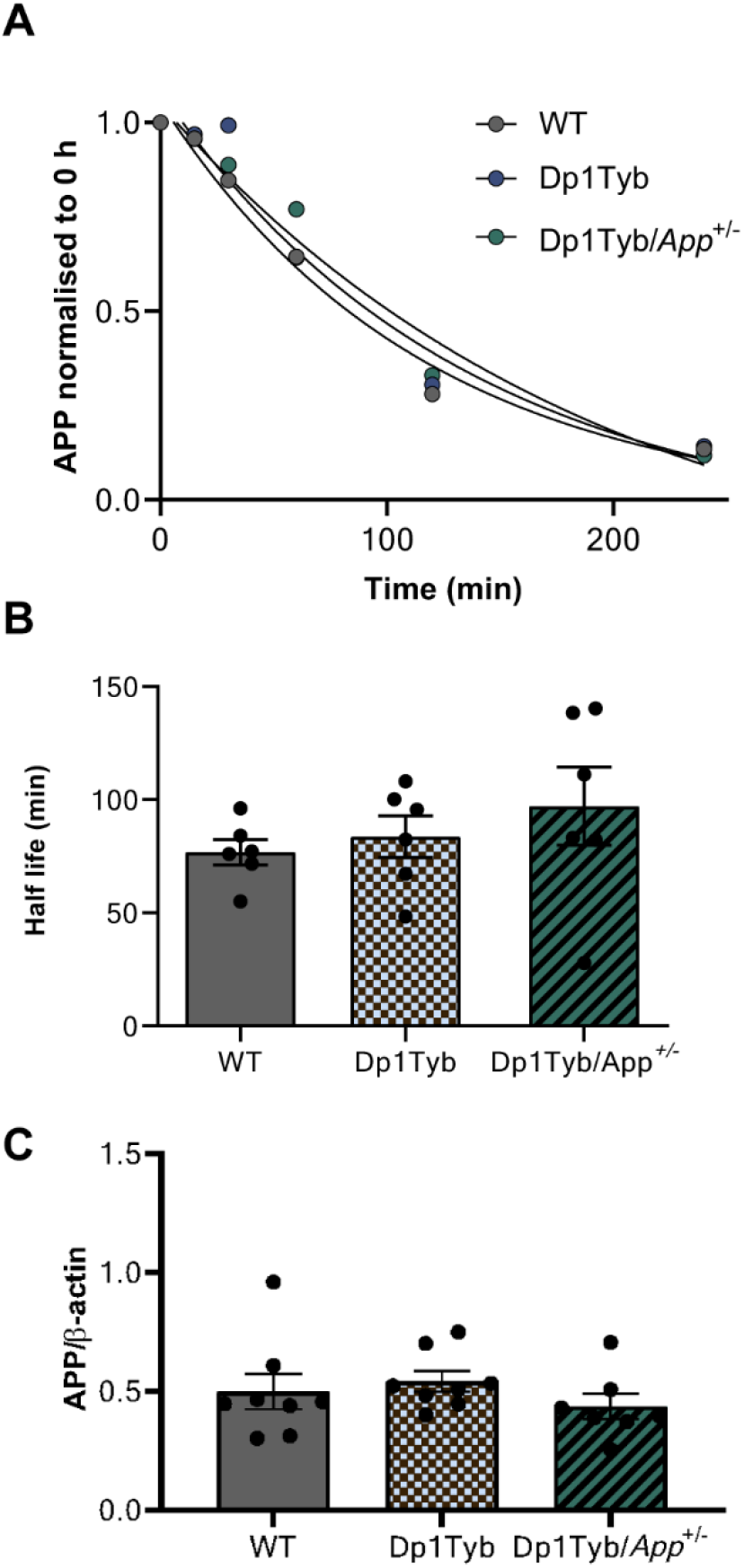
Trisomy of Hsa21-homologous genes including or excluding App does not affect APP half-life in MEFs. **A)** Degradation curve of APP in Dp1Tyb, Dp1Tyb/*App*^*+/*-^ and WT MEFs. **B)** Half-life of APP is not significantly different in Dp1Tyb, Dp1Tyb/*App*^*+/*-^ and WT MEFs (One-way ANOVA, *p* = 0.48, N = 5/6). Average APP half-life in minutes: Dp1Tyb = 84 ± 9; Dp1Tyb/*App*^*+/-*^ = 97 ± 17; WT = 77 ± 6). **C)** APP abundance is not significantly different in Dp1Tyb, Dp1Tyb/*App*^*+/*-^ and WT MEFs (One-way ANOVA, *p* = 0.77, N = 6). Average APP/β-actin: Dp1Tyb = 0.93 ± 0.07; Dp1Tyb/*App*^*+/-*^ = 0.83 ± 0.1; WT = 0.93 ± 0.15). Each dot corresponds to a biological repeat (i.e. an independent MEF line used). For each biological repeat, three technical repeats (i.e. western blot) were performed. Error bars = SEM. All full uncropped western blots are available at Figshare.

### Three-copies of Hsa21 mouse homologues in the Dp1Tyb region do not alter amyloid-β production or peptide ratios

Trisomy of genes on Hsa21 other than *App* can modulate the ratio of amyloid-β in vivo [2]. Levels of amyloid-β produced from the endogenous *App* gene were below the limit of detection in MEF culture media; thus, to determine if peptide ratios were altered we transfected MEFs with a βCTF-3xFLAG plasmid to overexpress APP-β-CTF and we quantified amyloid-β. The absolute concentrations of amyloid-β_40_ and amyloid-β_42_ and the ratio of the two peptides were not altered (N = 3) (**Fig 4A-C**). amyloid-β_38_ levels were below the limit of detection and were not analysed.

**Figure 4.**
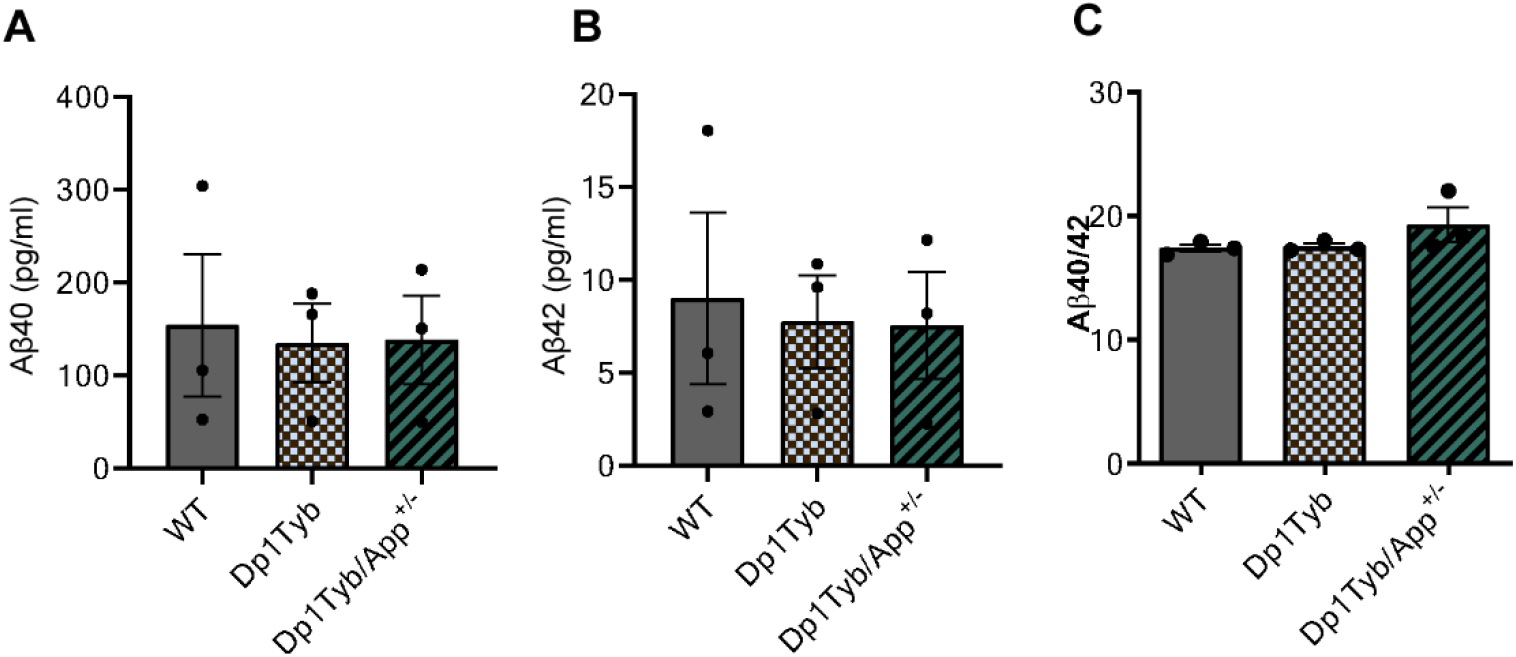
Trisomy of Hsa21-homologous genes, including or excluding App, does not affect Aβ40/Aβ42 ratio. In WT, Dp1Tyb and Dp1Tyb/*App*^*+/-*^ MEFs overexpressing APP-β-CTF led to no difference in **A)** Aβ40 abundance (Dp1Tyb = 134.9 ± 42.5; Dp1Tyb/*App*^*+/-*^ = 138.2 ± 47.75; WT = 154.1 ± 76.59. One-way ANOVA, *p* = 0.97, N = 3); **B)** Aβ42 abundance (Dp1Tyb = 7.78 ± 2.5; Dp1Tyb/*App*^*+/-*^ = 7.56 ± 2.88; WT = 9.02 ± 4.61 in pg/ml. One-way ANOVA, *p* = 0.95, N = 3) or **C)** Aβ40/Aβ42 ratio (Dp1Tyb = 17.51 ± 0.24; Dp1Tyb/*App*^*+/-*^ = 19.29 ± 1.37; WT = 17.38 ± 0.31. One-way ANOVA, *p* = 0.26, N = 3). Each dot corresponds to a biological repeat using an independent MEF lines. Error bars = SEM.

## Discussion

Here we compared the biology of early endosomes and APP in MEFs isolated from the Dp1Tyb and Dp2Tyb mouse models to determine if this system can be used to investigate the Hsa21 genes responsible for the changes to early endosomes and APP biology that occur in DS. Moreover, the workflow described here may be useful for the systematic quantification of RAB5^+^ endosome size and number in other cellular models as an alternative to the use of electron microscopy. We found that this DS MEF model did not recapitulate endosomal enlargement, likely because of the dosage insensitivity of *App* in this system. This is consistent with a previous report that showed raised levels of the *App* gene product β-CTF are necessary for DS-associated endosomal enlargement [40].

MEFs are embryonic peripheral cells and further changes to biology may be observed in neuronal cells or in the context of aging. However, cellular dysfunction including in the endo-lysosomal system occurs in iPSCs, organoids, fibroblasts and lymphoblastoid cells isolated from individuals with DS [14,16,23,41]. Future research could quantify APP expression in Dp1Tyb primary neurons to determine whether the lack of *App* dosage sensitivity in MEFs is a result of the embryonic origin of the cells or because of cell-type specific biology. Previous studies have been inconsistent on the dose sensitivity of *APP* in different tissues and models, suggesting that APP production is tightly regulated [42–44]. Since three copies of *APP* are sufficient for AD development and APP is the precursor of amyloid-β [1], studying the regulation of APP expression in different tissues and over time could be pivotal to gain further understanding of AD.

To further investigate APP processing *in vitro* we determined the ratio of amyloid-β_40_ and amyloid-β_42_ peptides and their absolute abundances in MEFs transfected with human β-CTF. The amyloid-β_40/_amyloid-β_42_ ratio was not altered in Dp1Tyb or Dp1Tyb/*App*^*+/-*^ MEFs compared to WT controls. This suggests that the additional copy of genes in this region is not sufficient to modulate the processing of APP-CTF to form amyloid-β in fibroblasts. Alić et al. (2020) observed that organoids trisomic for Hsa21 also failed to show an alteration in amyloid-β_40/_amyloid-β_42_ ratio, but the authors observed an increase in the absolute concentration of amyloid-β_40_ and amyloid-β_42_ produced, together with an increase in total APP which we did not observe in our mouse derived model system. Future research could use brain tissue from Dp1Tyb and Dp2Tyb mice to verify that APP, amyloid-β_40_ and amyloid-β_42_ abundance is increased in the mouse model and the lack of dosage sensitivity is a feature of MEFs. Use of brain tissue at different time points could enable investigation of the progressive changes over life-span, which cannot be investigated using primary cells. In addition, both Dp1Tyb and Dp2Tyb mouse models contain three copies of a number of mouse homologous chromosome 21 genes, which make them more physiologically relevant than single-gene transgenic models [45].

In conclusion, alternative models to the MEF system investigated here are required to understand how additional copies of genes on Hsa21 change endo-lysosomal and APP biology. These biological processes are proposed to underlie the early development of AD in people who have DS and the identification of alternative model systems will further understanding of this important research area.

## Material and methods

### Mouse breeding and husbandry

This study was conducted in accordance with ARRIVE2.0 [46]. The mice involved in this study were housed in controlled conditions in accordance with Medical Research Council guidance (*Responsibility in the Use of Animals for Medical Research*, 1993), and experiments were approved by the Local Ethical Review panel (MRC Prion Unit, University College London) and conducted under License from the UK Home Office, according to the revised Animals (Scientific Procedures) Act 1986.

Cage groups and genotypes were pseudo-randomised, with a minimum of two mice and a maximum of five in each cage; groups were weaned with members of the same sex. Mouse houses, bedding and wood chips, and continual access to water were available to all mice, with RM1 and RM3 chow (Special Diet Services, UK) provided to breeding and stock mice, respectively. The water provided was reversed osmosis (RO water). Cages were individually ventilated in a specific pathogen-free facility. Mouse used to generate the breeding stock for this study were euthanised by exposure to a rising concentration of CO_2_ gas followed by confirmation of death by dislocation of the neck, according to the revised Animals (Scientific Procedures) Act 1986. The animal facility was maintained at a constant temperature of 19-23°C with 55 ± 10 % humidity in a 12 h light/dark cycle.

Dp(16Lipi-Dbtb21)1TybEmcf [Dp1Tyb] (MGI:5703853) and Dp(16Mis18aRunx1)2TybEmcf [Dp2Tyb] (MGI:5703854) mice were imported from the Francis Crick Institute and colonies were maintained by backcrossing to C56BL/6J. B6.129S7-*App*^*tm1Dbo*^/J [*App*^*+/-*^] (MGI:2136847) mice were imported from the Jackson Laboratory and the colony was maintained by crossing heterozygous knockouts with C57BL/6J animals. To generate progeny for the MEFs used in this project Dp1Tyb mice were crossed with *App*^+/-^ or C57BL/6J mice; Dp2Tyb mice were maintained by crossing with C57BL/6J animals. Dp1Tyb, Dp2Tyb, *App*^+/-^ colonies were fully inbred for >10 generations on the C57BL/6J genetic background.

### Mouse embryonic fibroblasts (MEFs)

Mouse Embryonic Fibroblasts were generated from timed matings; at E14 pregnant females and embryos were culled by a schedule one method. Briefly, the pregnant female mouse in the mating was euthanized, and dissection for the collected embryos was carried out under sterile condition in a laminar flow hood. The uterine horn was dissected and rinsed in 70 % ethanol (v/v) and placed into a 100 mm Petri dish. Each embryo was separated from its placenta and embryonic sac. The embryo was decapitated and the head and body were transferred to a 1.5 ml Eppendorf tube containing PBS and delivered for genotyping (heads) and MEF generation (bodies). Red organs were removed from embryo bodies and remaining tissue was minced with % trypsin-EDTA prior to dissociation by pipetting, cells were isolated by centrifugation and plated on 0.1 % gelatin-coated plates in DMEM + GlutaMax, 10 % FBS and 1 % Penicillin-streptomycin (culture at 37°C in 5 % CO_2_).

### Genotyping

Genotyping of Dp1Tyb, Dp2Tyb, *App*^+/-^, and Dp1Tyb/*App*^+/-^ mice was outsourced to TransnetYX (Cordova TN, USA) using a proprietary qPCR-based system.

### Generation of the βCTF-3xfLAG plasmid, the GFP-Rab5CA plasmid and nucleofection

Briefly, the β-CTF sequence was amplified from a βCTF-EGFP plasmid (kind gift of Dr Jiang (New York University, USA), then ligated into a pCI-Neo vector. Then the APP signal peptide sequence was ligated into the 5’ region and the 3xFLAG into the 3’ region. GFP-Rab5CA (Q79L) (RAB5CA), was a kind gift from Sergio Grinstein sourced from Addgene (Addgene plasmid # 35140 ; http://n2t.net/addgene:35140 ; RRID:Addgene_35140 [37]). These plasmids were transfected into TOP10 competent cells under ampicillin selection and DNA was prepared from cultures with a QIAprep Spin Miniprep Kit (QIAGEN) according to manufacturer’s instructions. An Amaxa Nucleofector 2b Device and a Mouse Embryonic Fibroblast Nucleofector Kit 1 (Lonza) were used to transfect MEFs with βCTF-3xFLAG plasmid using program N-024 of the Nucleofector (**Supplementary Figure 3**).

### Cycloheximide pulse chase

13 h after plating, MEF media was changed and cycloheximide solution (30 μg/ml per well) or ddH_2_O (negative control) were added. Cells were collected at 6 timepoints from cycloheximide addition: 0 h, 15 min, 30 min, 1 h, 2 h, 4 h in ice-cold RIPA buffer (150 mM sodium chloride, 50 mM Trizma hydrochloride, 1 % NP-40, 0.5 % sodium deoxycholate, 0.1 % SDS) + 1:100 protease inhibitor (Protease inhibitor cocktail I). The cell suspension was centrifuged for 15 min at 24 000 rcf at 4°C. APP abundance at each timepoint was normalized to the value at time 0 h. Half-life was calculated using the One Phase Decay (nonlinear regression) function on GraphPad Prism. The values obtained for each technical repeat (i.e. gel) were averaged together to obtain one half-life value per genotype per experimental repeat, such that independent biological replicates were used as the experimental unit. These values were then compared with a one-way ANOVA test on GraphPad Prism.

### Aβ peptides measure

The Mesoscale amyloid-β 6E10 Triplex Assay (Meso Scale Discovery, MSD) was used to determine the concentration of amyloid-β isoforms (amyloid-β_38_, amyloid-β_40_, amyloid-β_42_) in media collected from MEFs and diluted 1:2 in Dilutor 35. A MESO SECTOR S 600 plate reader (MSD) was used to read the plate.

### Western blotting

Pierce 660nm Protein Assay Reagent was used to measure protein concentrations using a standard of Bovine Serum Albumin (BSA) in PBS (3000 – 0 μg/ml). Samples were denatured in NuPAGE LDS 4X and 2-mercaptoethanol by boiling at 95°C for 5 min. Bolt 4-12 % Bis-Tris Plus Gels and Bolt MES SDS Running Buffer 20X were used for protein separation before transfer to nitrocellulose membranes (Transblot Turbo Transfer Pack, Bio-Rad) using a Transblot Turbo 0.2 μm (Bio-Rad). Proteins were blocked in 5 % skimmed milk in PBS prior to incubation with primary antibody (anti-APP A8717 1:5000 Sigma Aldrich) at 4°C overnight prior to incubation with anti-rabbit HRP. Membranes were developed using Super Signal West Pico Chemiluminescent Substrate. ImageJ was used to quantify the signal from bands and the linearity of APP signal was confirmed by western blot of endogenous APP (doubling-dilutions).

### Immunocytochemistry

Cells were washed in PBS then fixed in 4 % PFA for 20 min prior to permeabilization with 0.05 % saponin/PBS for 10 min. Cells were blocked with 5 % BSA/PBS for 1 h before overnight incubation with primary antibodies in 1 % BSA/PBS (RAB5 21435 1:200 Cell Signalling and anti-Integrin-β1 MAB1997 1:1000, Millipore) at 4°C prior to washing and incubation with secondary antibodies (anti-rabbit AlexaFluor-546 [A11-35] and anti-mouse AlexaFluor-633 [A21052] Thermofisher) in 1 % BSA/PBS. Cells were mounted on SuperFrost adhesion slides (VWR International) with ProlongGold + DAPI.

### Imaging

Images were taken on Confocal microscopes Zeiss Observer LSM800 or Zeiss Examiner LSM880. Each image was taken with a 63×1.4 Oil Plan Apochromat objective in two channels. Z-stacks at 150 nm interval between slices were taken to include the whole cell. Pixel size was equal to x, y = 0.05 μm, z = 0.15 μm. The pinhole size was equal to 1 Airy Unit of the 546 channel. Deconvolution for RAB5 signal was performed with Huygens software signal/noise ratio = 15. ImageJ software was used to clear the space surrounding the cells and to measure their volume. Briefly, the surface of the cell was smoothed and thresholded in 3D; everything outside the cell was cleared, using a custom macro (supplementary Fig 4). Imaris software was used to build a 3D reconstruction of the staining after deconvolution. Objects were identified using the surfaces function, with smoothing disabled and thresholding with background subtraction using default settings. This allowed us to make an accurate measurement of a large number of endosomes in three dimensions. Volume data were generated by the software and imported in excel. Endosomal volume (μm^3^) was used to calculate endosomal size. The size parameters of endosomes between the 50 and 90 percentiles were determined in WT MEFs transfected with PBS, and this information was used to classified endosomes in small (0-50 percentile), medium (50-90 percentile) and large (90-100 percentile) bins. A nested ANOVA was used to compare the size of large endosomes in MEFs transfected with PBS vs RAB5.

### Experimental Design and Statistical analysis

Sample size was determined with either a power calculation using pilot data (Dp1Tyb) or based on sample availability (Dp2Tyb). Sample order in all experiments (including during culture, western blotting and MSD assay) was randomized but balanced by genotype. All experiments and data analysis undertaken blind to genotype. All statistical tests were performed with IBM SPSS Statistics Version 2.5 and GraphPad Prism Version 8.4.2. All data is reported as mean ± SEM. All data was checked for normality of distribution and homogeneity of samples; sample distribution was tested with a Levene’s test, and data normality was tested with a Kolmogorov-Smirnov test. If the assumptions of normality and homogeneity of variance were verified, parametric tests were used to analyse data; otherwise non-parametric tests were used. For each experiment, the effect of genotype and sex was assessed using a multivariate ANOVA test. If the effect of one or more of the variables was significant, the variable was tested separately using ANOVA test, t-test or their non-parametric equivalents.

## Acknowledgements and Funding

C.C. was funded by an Alzheimer’s Society PhD studentship (AS-PhD-16-003 2017-20) awarded to F.K.W and E.M.C.F. F.K.W holds an Alzheimer’s Research UK Senior Research Fellowship (ARUK-SRF2018A-001). F.K.W. is also supported by the UK Dementia Research Institute (UKRI-1014) which receives its funding from DRI Ltd, funded by the UK Medical Research Council, Alzheimer’s Society and Alzheimer’s Research UK. F.K.W. and E.M.C.F. also received funding that contributed to the work in this paper from the MRC via CoEN award MR/S005145/1. E.M.C.F. received funding from a Wellcome Trust Strategic Award (grant number: 098330/Z/12/Z) awarded to The London Down Syndrome (LonDownS) Consortium and a Wellcome Trust Joint Senior Investigators Award (grant numbers: 098328, 098327). DM is supported by NIHR GOSH BRC award 17DD08. The funders had no role in study design, data collection and analysis, decision to publish, or preparation of the manuscript. For the purpose of Open Access, the author has applied a CC-BY public copyright licence to any Author Accepted Manuscript version arising from this submission.

The LonDownS Consortium comprises Andre Strydom^1,2^, Elizabeth M.C. Fisher^3^, Frances K. Wiseman^4^, Dean Nizetic^5,6^, John Hardy^4,7^, Victor L. J. Tybulewicz^8,9^ and Annette Karmiloff-Smith^10. 1^Department of Forensic and Neurodevelopmental Sciences, Institute of Psychiatry, Psychology and Neuroscience, King’s College London, London, UK. ^2^Division of Psychiatry, University College London, London, UK. ^3^Department of Neuromuscular Diseases, Queen Square Institute of Neurology, University College London, Queen Square, London, UK. ^4^The UK Dementia Research Institute, University College London, Queen Square, London, UK. ^5^Blizard Institute, Barts and the London School of Medicine, Queen Mary University of London, London, UK. ^6^Lee Kong Chian School of Medicine, Nanyang Technological University, Singapore, Singapore. ^7^Reta Lila Weston Institute, Institute of Neurology, University College London, London, London, UK. ^8^The Francis Crick Institute, London, UK. ^9^Department of Immunology and Inflammation, Imperial College, London, UK. ^10^Birkbeck University, London, UK.

We thank Amanda Heslegrave (UCL-DRI) for assistance with this project.

F.K.W. has undertaken consultancy for Elkington and Fife Patent Lawyers unrelated to the work in the manuscript and is also a PLoS One Academic Editor. This does not alter our adherence to PLoS One policies on sharing data or materials.

## Figure legends

**Supplementary Figure 1.**
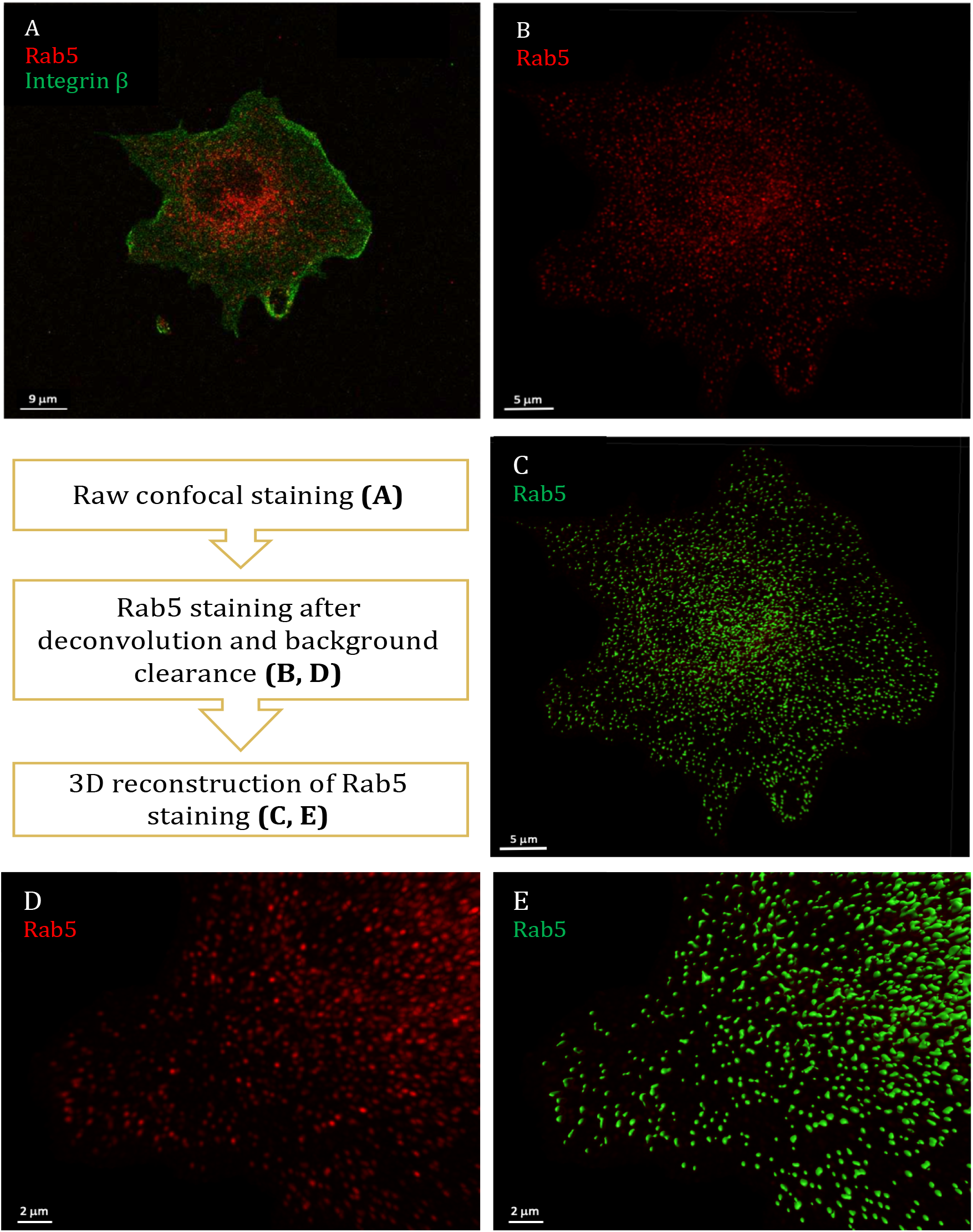
Process of quantification of RAB5^+^ endosomal staining. **A)** WT MEF stained for Integrinβ (cell membrane, green) and RAB5 (endosomes, red). **B, D)** Endosomal staining after deconvolution and background clearance **C, E)** 3D reconstruction of endosomal staining. Deconvolution and 3D reconstruction to accurately quantify the volume of endosomes. Z-stacks of each cell were taken with 150 nm interval between slices and fixed voxel volume (x = 50 nm, y = 50 nm, z = 150 nm) on confocal microscopes LSM800 or LSM880. Each stack was deconvolved using Huygens software to improve image signal to noise and resolution. ImageJ software was used to remove the background with a macro written by Dr Dale Moulding. Imaris software was used to reconstruct the deconvolved staining in 3D. The area of Integrinβ was used to create a mask to define cellular volume.

**Supplementary Figure 2.**
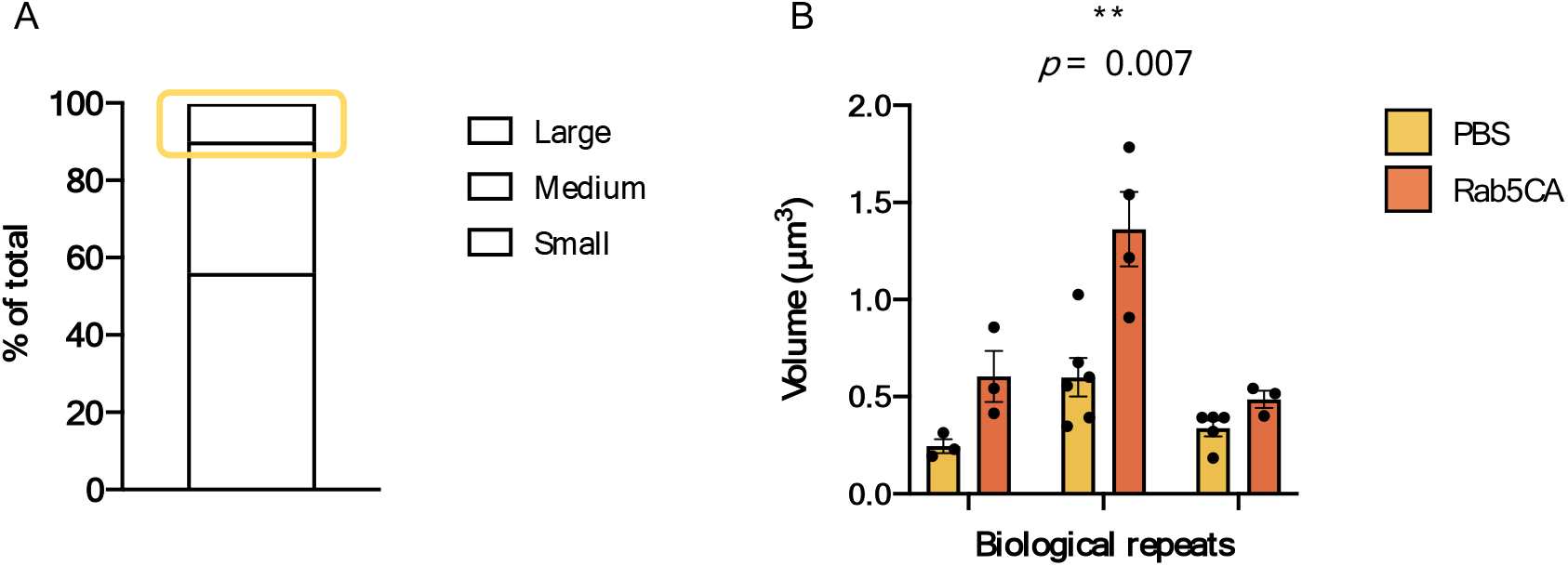
Distribution and quantification of endosomes in MEFs transfected with PBS and RAB5CA. **A)** The normal distribution of endosomal size in WT MEFs transfected with PBS was determined to define the parameters for classification of “large” endosomes (small: endosomes in the 0 – 50 percentile, medium: endosomes in the 50 – 90 percentile, large: endosomes in the > 90 percentile). **B)** A nested t-test showed that ‘large’ endosomes in cells transfected with RAB5CA had a significantly higher volume than the endosomes in cells transfected with PBS (*p = 0*.*007*, N = 3 of biological repeats). *The dots indicate the average volume of the ‘large’ endosomes in one cell imaged (technical repeat). Error bars = SEM*.

**Supplementary Figure 3.**
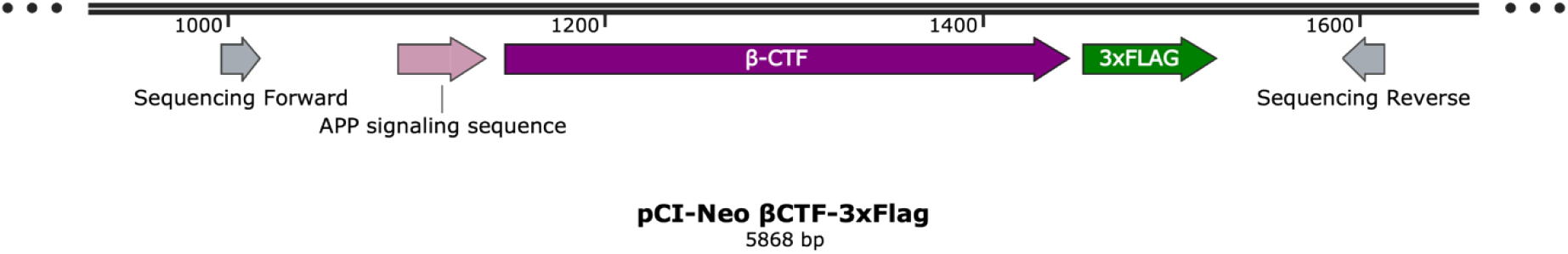
Detail of the pCI-neo βCTF-3xFLAG plasmid map. The APP signalling sequence was inserted in a pCI-neo plasmid followed by the β-CTF fragment of APP and by a 3xFLAG sequence. The primers used for sequencing the insert (*sequencing forward and reverse*) are also shown.

**Supplementary Figure 4.**
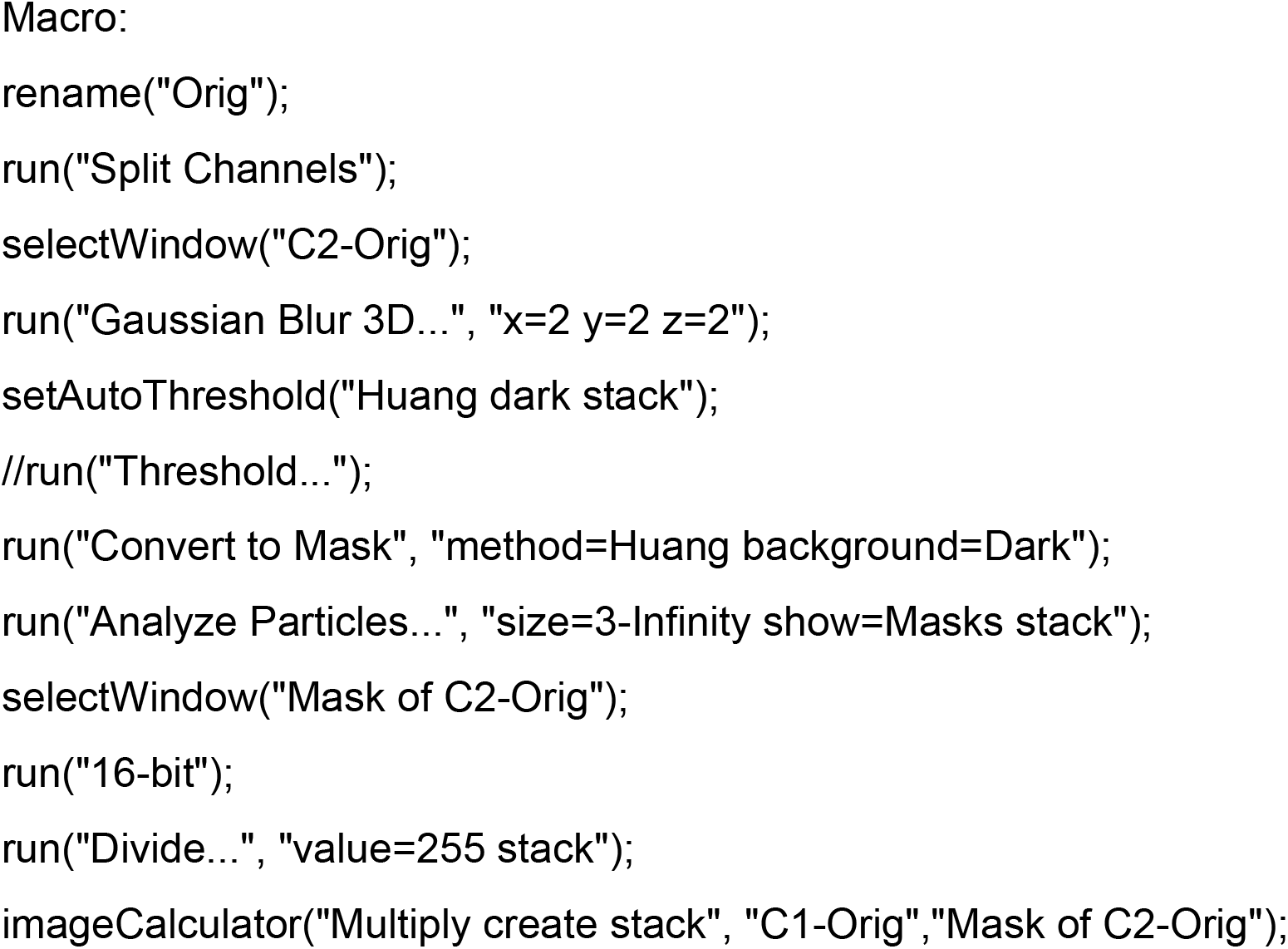
Custom ImageJ Macro. Macro designed by Dr Dale Moulding to smooth the cell surface and clear its outside in 3D, enabling accurate quantification of the volume of the cell and of the number and volume of endosomes.

